# Slow-varying normalization explains auditory steady-state masking interactions in human EEG

**DOI:** 10.1101/2025.08.25.672175

**Authors:** Divya Gulati, Supratim Ray

## Abstract

The inherent ability of sensory neurons to entrain to modulations in the temporal structure of an auditory stimulus gives rise to the auditory steady-state response (ASSR). Simultaneous presentation of multiple acoustic stimuli by frequency tagging them to generate ASSRs at different frequencies is routinely employed for hearing threshold determination and cognitive studies. However, the nature of ASSR interactions as a function of competing modulation frequencies in the absence of overt behaviour and the underlying neural mechanisms have not been well studied. Such interactions have previously been studied with visual stimuli that generate steady-state visually evoked potentials (SSVEPs) and explained using a normalization model. Here, we tested whether similar interactions are observed in the auditory domain, and if so, can be explained using a similar model. We played sinusoidally amplitude-modulated stimuli simultaneously modulated at two frequencies while we recorded a 64-channel EEG from subjects who passively listened to these sounds with closed eyes. We used multiple modulation frequencies and depths to characterize the ASSR modulation masking response profile. We observed that modulation frequencies closer to each other suppress the ASSR strongly than frequencies that are farther apart, similar to the interaction observed in the visual domain using SSVEPs. The observed suppression was captured by a slow-varying normalization model, which was initially used to explain SSVEP interactions. We obtained a band-pass shaped suppression profile with a low-pass cutoff that matched to that observed for SSVEP interactions. Our findings highlight the universality of the normalization model in accounting for masking interactions across modalities.

## Introduction

Neurons in the auditory cortex often phase-lock to rhythmically modulated acoustic stimuli. This frequency-following time-locked electrophysiological response is known as the Auditory steady-state response (ASSR) (Galambos et al., 1981; Kuwada et al., 1986). ASSRs are robust, easy to record, and can be measured non-invasively using electro- or magneto-encephalography (EEG or MEG) (Kuwada et al., 1986; Ross et al., 2000). They are an ideal candidate to track cortical responses by ‘frequency tagging’ each stimulus with a different frequency (Fujiki et al., 2002). Consequently, ASSRs are widely used for studying auditory selective attention (Lazzouni et al., 2010; N. Müller et al., 2009), building ASSR-based brain computer interface (BCI) (Lopez et al., 2009), and evaluating hearing thresholds, especially in infants, the elderly, and in patients who are unable to provide verbal feedback (Ishida & Stapells, 2012; John et al., 1998; Lins & Picton, 1995; Sek et al., 2015).

Such studies often employ simultaneous presentation of multiple spectrally overlapping stimuli, which leads to non-linear interactions among the concurrently evoked ASSRs irrespective of any behaviour. Previous studies have noted a reduction in amplitude of ASSRs when two sinusoidally amplitude-modulated (SAM) stimuli at different modulation frequencies were concurrently presented (Draganova et al., 2002; Ishida & Stapells, 2012; John et al., 1998; Lins et al., 1995; Lins & Picton, 1995; Ross et al., 2003). This interaction depends on factors such as carrier frequency and intensity of SAM. However, its dependence on the modulation frequency remains underexplored. An EEG study observed no effect of modulation frequency of SAMs on ASSR amplitude (Gransier et al., 2017), contrasting the modulation frequency-dependent masking observed in psychoacoustic studies (Bacon & Grantham, 1989; Grantham & Bacon, 1991; Sek et al., 2015). Due to these conflicting differences, whether masking is a function of modulation frequency remains unclear. Additionally, the mechanism governing this interaction is also not well understood.

In the visual modality, analogous to ASSRs, attention and BCI studies have used steady-state visually evoked responses (SSVEPs), which are produced by flickering or counterphasing grating stimuli (Ding et al., 2006; M. M. Müller et al., 2006; Vialatte et al., 2010). Interactions between multiple SSVEPs have been extensively studied and recently been explained using gain control or normalization models (Candy et al., 2001; Gulati & Ray, 2025; Liza & Ray, 2022; Salelkar & Ray, 2020; Tsai et al., 2012). SSVEP masking interactions were found to depend on the orientation and temporal frequency difference between the two simultaneously presented gratings. Parallel gratings at nearby frequencies caused the strongest SSVEP suppression, which was reduced as the difference between the two frequencies increased, termed as “target-frequency dependent” suppression (Gulati & Ray, 2025). This was accounted for by a slow-varying divisive normalization model, whereby the input drive to a neuron is “normalized” divisively by a low-pass filtered weighted sum of the activity of the neuronal neighbourhood.

In the auditory domain, the normalization model has only been utilized to explain spectro-temporal contrast gain control against statistically stationary background stimuli in primary auditory cortex neurons of mice and ferrets (Cooke et al., 2018; Rabinowitz et al., 2011). The latter study also extended the model to capture the dependence of neuronal gain on spectral overlap between two simultaneously presented dynamic random chord stimuli. Therefore, we speculated whether the normalization model could account for modulation masking in ASSRs and whether computations employed by the neuronal pool for temporal integration follow canonical principles akin to those found in the visual modality. Since sensory systems across modalities share similar architecture (King & Nelken, 2009), we tested whether ASSRs interact in ways comparable to SSVEPs, and if so, whether a slow-varying normalization model can explain these interactions. To empirically address this, we presented tones that were amplitude-modulated at two modulation frequencies simultaneously at various modulation depths through loudspeakers while recording EEG from human subjects.

## Methods

### Participants

Twenty-four healthy adults (mean± std of 26.91±3.84 years, range 22-38 years, 12 females) were recruited from the Indian Institute of Science community. All the subjects reported having normal hearing levels with no abnormalities. All subjects but one were right-handed. A written consent was obtained from all the subjects after explaining the experiment paradigm. Participation in the task was voluntary, and subjects were monetarily compensated. All the experiments were performed in accordance with the protocol approved by the Institutional Human Ethics Committee of the Indian Institute of Science, Bangalore.

### EEG Setup and Data Acquisition

A 64-channel active electrode (actiCap) EEG system (BrainAmp DC EEG acquisition system, Brain Products GmbH) was used to record the measurements. All the electrodes were placed on the subject’s head using a cap that followed the international 10-10 system, with FCz as the reference electrode. EEG signals were sampled at 1000Hz and were band-pass filtered between 0.016Hz (first-order filter) and 250Hz (fifth-order Butterworth filter). Signals were digitized at 16-bit resolution (0.1µV/bit). Across the entire recording duration for each subject, electrode impedance was kept below 25kΩ, with the average impedance of the final set of electrodes being 6.92±2.97 KΩ (mean± std).

### Experimental Paradigm and Stimuli

The recording was conducted in a quiet room to minimize irrelevant sounds. Participants sat in a dark room with their eyes closed and were instructed to listen to auditory sounds passively. They were asked to maintain the same posture while keeping their eyes closed to minimize movement and blink artefacts. 1-2 minute breaks were given to participants after every 5-6 minutes to maintain a constant arousal state and prevent them from falling asleep. During the breaks, the investigator talked with the participants to keep them in an alert state. Sound stimuli were generated at a sampling rate of 44100Hz using a custom-written MATLAB (The MathWorks, Inc.; <u>RRID: SCR_001622</u>) script and were presented using NIMH Monkeylogic software toolbox in MATLAB. Stimulus onset markers were recorded using a digital I/O card (USB 6210, National Instruments) in the EEG data file. Stimuli were played at 75-80dB SPL using a multidirectional speaker (Marshall Kilburn II).

Each participant came in for a session lasting 3-3.5 hours. Each session had two experiments, which were run in counterbalanced order. The first 10 participants also did additional auditory and visual protocols, as reported in our previous study (Gulati & Ray, 2024).

The first experiment consisted of sinusoidally amplitude-modulated (SAM) sounds generated using the following equation:

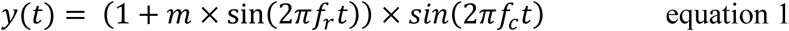

Where *m* is the modulation depth, 𝑓_𝑟_ is the modulation frequency in Hz, *f_c_* is the carrier frequency in Hz, and t is the time in seconds. The stimuli had a carrier frequency of 1kHz and were amplitude-modulated at a modulation frequency of 22Hz to 58Hz, in steps of 6Hz and at a modulation depth of 6.25% to 100% (5 intervals spaced on a log scale).

In the second experiment, a sinusoidally amplitude-modulated carrier was modulated at two different modulation frequencies. Here, two modulating waveforms were added prior to a single multiplication with the carrier waveform (as we intended to keep the carrier frequency the same for both). The stimuli were generated using the following equation:

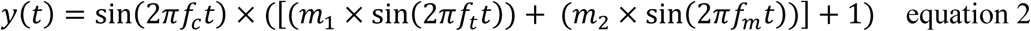

where, 𝑓_𝑡_is the target modulation frequency, and 𝑓_𝑚_is the mask modulation frequency. The stimuli had a carrier frequency (*f_c_*) of 1kHz. The modulation frequency for the target was set at 40Hz, whereas the mask could take any modulation frequency between 22-58Hz (in steps of 6Hz). The modulation depth (*m*_1_ and *m*_2_) could take one of the three values: 0, 25% or 50%. When the mask has a modulation depth of 0% (i.e., *m*_2_ = 0), that condition is referred to as the ‘target’ only condition or single SAM. Other conditions are called a dual SAM, equivalent to simultaneously presenting two amplitude-modulated sinusoids. Previous studies have used comparable stimuli, albeit with differences in carrier type, phase differences between mask and target components, and the normalization constant (Bacon & Grantham, 1989; Slugocki et al., 2017). Here, we chose this formulation to keep our stimuli as consistent as possible with our previous SSVEP studies.

In both experiments, each stimulus was 800ms long (including a rising and falling off time of 10ms each), followed by a 200ms inter-stimulus interval. An additional 800ms stimulus, with zero carrier amplitude, was also presented. In this condition, no audio was perceived. We used this condition as a baseline for experiment 1. For experiment 2, we used a 1kHz unmodulated pure tone, i.e., the condition where the modulation depth for both target and mask stimuli was kept at 0% as the baseline condition. To keep the loudness across the sounds the same, we normalized the sounds by the maximum amplitude across all the sounds for that experiment. We obtained 36 and 64 stimulus conditions for the first and second experiments, and each stimulus was repeated 40 times in a pseudorandom order. Here, each stimulus repeat is referred to as a trial.

### EEG Artefact Rejection

We used an automated pipeline to get artefact-free data (See Gulati & Ray, 2024; Murty & Ray, 2022). In brief, to obtain bad trials across electrodes, we applied thresholding to the root mean squared (RMS) value of time-domain waveforms (upper RMS cutoff - 35µV and lower RMS cutoff - 1.5µV), followed by thresholding of multitapered power spectral density (PSD; calculated using Chronux toolbox, version 2.10; Bokil et al., 2010; <u>RRID: SCR_005547</u>; available at http://chronux.org) (between 200ms before the stimulus onset and 1200ms post-onset, 0-200Hz) for each trial (any trial for which the PSD at any frequency was beyond the mean +-6 SD was considered as bad). Subsequently, we rejected those electrodes that had more than 30% of all trials labelled as bad. Additionally, if any trial was bad in more than 10% of all the remaining good electrodes, it was labelled as a common bad trial. Further, if any trial was marked as bad in the subsets of the temporal, parieto-occipital and occipital electrodes (TP7, TP8, TP9, TP10, Pz, P1, P2, P3, P4, P5, P6, P7, P8, POz, PO3, PO4, PO7, PO8), it was added to the set of common bad trials. We then discarded any electrode with a PSD slope in the range of 56-84Hz less than zero. We rejected ∼30% (Experiment 1-29.36 ± 10.94 %; Experiment 2 – 29.26 ± 9%) of the data collected for each of the experiments. Overall, this procedure yields a common set of good trials across electrodes for each participant. We then rejected any participant from analysis with less than 75% of electrodes labelled as good. This way, we rejected two subjects from the first experiment 1 and three subjects from the second.

### Data Analysis

All the analyses of this data were done using custom-written scripts in MATLAB 2021a. Violin plots were generated using MATLAB 2024b. The analysis was done with a 250ms-750ms period, giving a resolution of 2Hz. Amplitude spectra were calculated by taking the Fast Fourier Transform of the signal averaged across all good repeats for a particular stimulus condition. The amplitude-modulated sinusoid evokes a strong ASSR at the modulation frequency of the sound. So, the amplitude difference was always calculated at the modulation frequency (for experiment 1- at 22, 28, 34, 40, 46, 52, and 58Hz and for experiment 2 at 40Hz, modulation frequency of target stimulus).

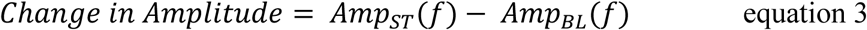

We computed the time-frequency spectrograms using the Chronux toolbox. We used the multitaper method using a single taper with a moving window size of 500ms and step size of 25ms, yielding a frequency resolution of 2Hz. We made two groups of electrodes corresponding to the left and right hemispheres, where a strong ASSR response was observed and were symmetrical across both the hemispheres (Right: TP8, TP10, CP6, P2, P4, P6, P8, PO4, PO8, O2; Left: TP7, TP9, CP5, P1, P3, P5, P7, PO3, PO7, O1). These sets of electrodes were used for final analysis.

We used topoplot.m function of the EEGLAB toolbox (Delorme and Makeig, 2004; RRID: SCR_007292) to generate the scalp maps.

#### Laterality Index

To measure if ASSR responses at 100% modulation depth are lateralized to one hemisphere, we calculated the Laterality Index (LI) –

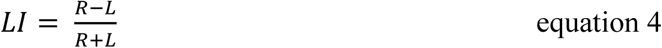

which corresponds to the difference in ASSR amplitude at the respective modulation frequency between right (R) and left hemisphere (L) electrodes, normalized by the sum of the responses of chosen electrodes in both hemispheres. The values of the laterality index range between -1 and +1, with -1 indicating complete left hemisphere dominance and +1 indicating right hemisphere dominance. Zero indicates a bilaterally symmetric response, i.e., no laterality.

#### Normalization model of auditory masking

In the auditory periphery, incoming sound signals undergo transformation from a pressure-time waveform to neural activity. The input target and mask SAMs are first spectrally analyzed by critical filterbanks located in the basilar membrane (Figure 1); these are linear gammatone filterbanks, with overlapping passbands, that half-wave rectify the sound and low-pass filter it (Chi et al., 2005; Dau et al., 1997a). This stage extracts the information about the carrier frequency and low-pass filtering essentially preserves the envelope of the signal. Since in our case the target and mask have the same carrier frequency, they will pass through the same filter. The filter outputs eventually reach the second stage, which represents the processing in the higher auditory regions, such as the primary auditory cortex. This stage consists of band-pass modulation filters. To model the processing of auditory steady-state response interactions through these filters, we used the optimal-tuned normalization model described previously (Gulati and Ray, 2025, equations 4 and 5) for steady-state visual evoked potential interactions. We kept the equations and the nomenclature unchanged to keep them comparable to the previous SSVEP study. The following equations were used:

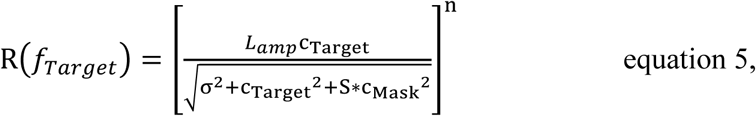

where R(*f_Target_*) is the change in amplitude at the modulation frequency of the target sound (at 40Hz). L_amp_ corresponds to the response of the target sound alone at the maximum modulation depth. c_Target_ and c_Mask_ represent the modulation depth of the target and mask components of the stimuli. σ and n represent the semi-saturation constant and exponent controlling the non-linear gain of the response, respectively. The numerator representing the excitatory drive gets its contribution only from the target once it crosses the modulation filter, because we were characterizing the responses at the target modulation frequency only. In the normalization signal, both target and mask interact non-linearly, represented by S, which is then scaled by the mask modulation depth. In the resulting inhibitory drive, the target and mask inputs can interact in two ways: inputs can be added linearly before or after they pass modulation filters, which often act like non-linear transducers. It is calculated as follows:

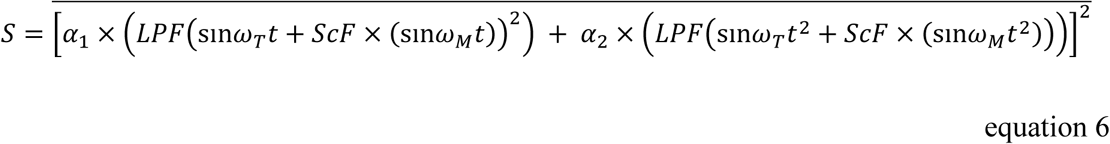

here, we assumed that the physical properties of the target and mask are transduced into neural responses, which are also changing sinusoidally, represented by ꞷ_T_ and ω_M_, target and mask frequencies (ꞷ equals to 2π*f*, where ‘*f*’ corresponds to the modulation frequency). After summing up the target and mask inputs before or after the exponentiation, we mean-corrected the signal to remove the DC component and passed it through a low-pass filter (LPF) to account for the temporal dynamics of the normalization drive. We used a fourth-order Butterworth filter. In the SSVEP study (Gulati & Ray, 2025), it was observed that the SSVEP amplitude suppression profile as a function of temporal frequency of mask grating (S values) was dependent on the orientation difference between the target and the mask grating. When gratings were parallel, a band-pass suppression profile was obtained, whereas for orthogonally oriented stimuli, a low-pass profile was obtained. For this reason, the model has *α_1_* and *α_2_*, which indicate the weights for two operations after passing through low-pass filters. When sinusoids, which are summed after exponentiation, pass through a low-pass filter, low-pass suppression values are obtained, as the value of ‘S’ would be higher for lower mask frequencies. When low-pass filtered inputs are first summed and then squared, the band-pass suppression profile is obtained. This can be understood by the trigonometric expansion of the terms:

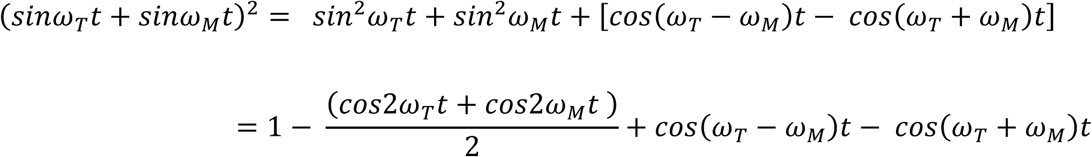

**Figure 1.**
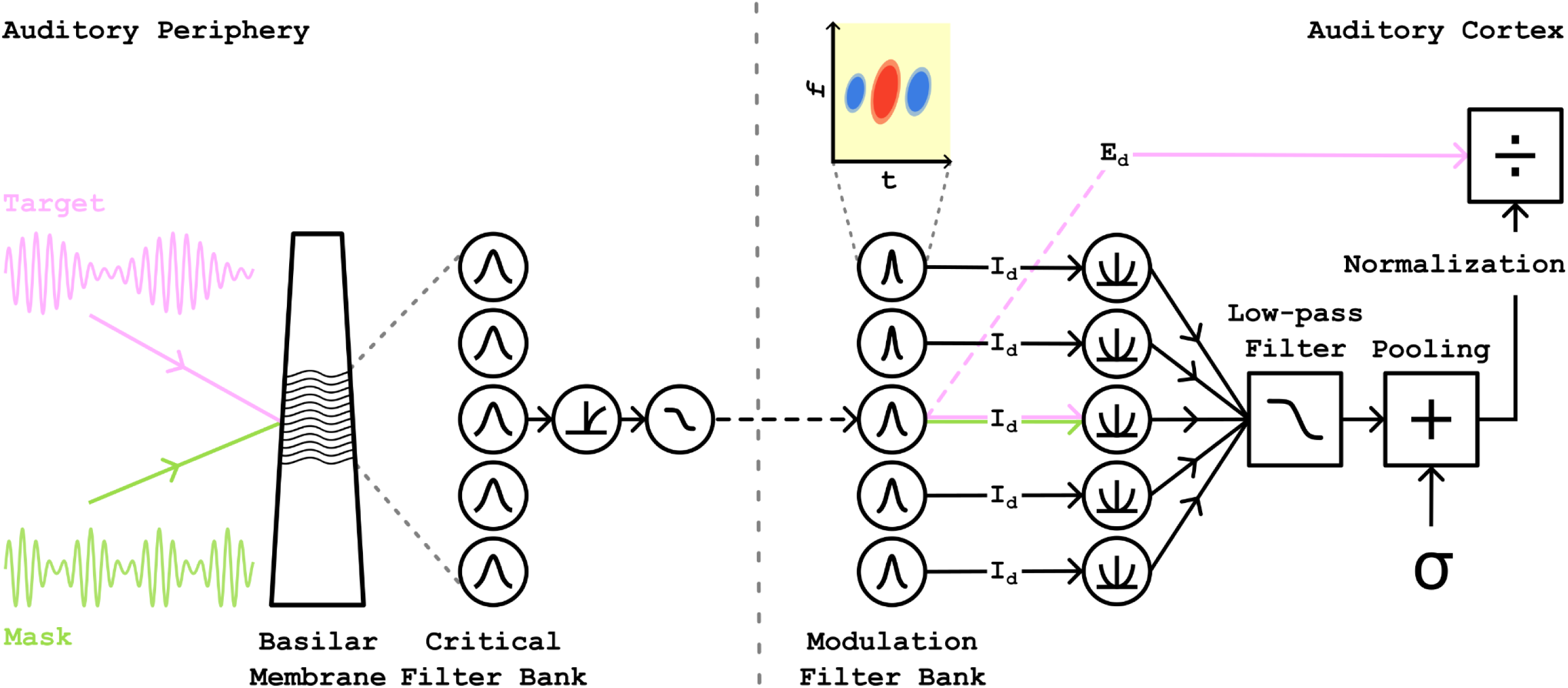
Schematic for auditory normalization model. The target and mask SAM stimuli first passes through the band-pass cochlear filters located at the basilar membrane. These filters are tuned for the carrier frequency of the SAM stimuli. The output of these filters is half-wave rectified and then passes through low-pass filter which passes on the information about the relatively slower modulation frequency of the stimulus. To keep the model simple and since ASSR interactions below 60Hz are primarily cortical in original, we bypassed the intermediary stages of neural processing. In cortex, the input first crosses through the modulation filter bank that tiles the modulation space. These modulation filters are spectro-temporal receptive fields of auditory neurons and analyse the modulation frequency of the stimulus. The bandwidth of these modulation filters is known to increase with increase in the centre frequency at which they respond. The excitation drive only consists of input from the target SAM (indicated by pink arrow). The inhibitory drive consists of input from both the target and mask SAM. When the modulation frequencies of two SAM lie within the bandwidth of the modulation filter bank, they are first summed and then passed through an expansive non-linearity. Otherwise, the two pass through different filters and are passed through expansive non-linearity separately and then summed. This drive is then pooled and passes through a low-pass filter, and then it scales the excitatory drive.

When this output is passed through a low-pass filter, maximum suppression will be obtained in two cases: if 𝜔_𝑀_ is small or if the difference between 𝜔_𝑇_ − 𝜔_𝑀_ is small, resulting in higher ‘S’ values for these cases.

Next, the inhibitory drive may not receive equal contributions from target and mask stimuli. Therefore, we added an additional parameter, ScF, which stands for scaling factor, which weighs the contribution of the mask stimulus in the inhibitory drive. S is not a function of time, as we averaged the value over time indicated by the bar above equation 6. The obtained value for S is then plugged back into equation 5 to get the response value. So, in this model, we fixed the value of n to 2, and we had six free parameters: L_amp_, constrained <15; σ, constrained <4; cutoff frequency of the filter, constrained <40; α_1_ and α_2,_ constrained >0; and *ScF*, constrained between 0.1 and 1.

We did the curve fitting using the fmincon function of MATLAB such that the sum of squared estimate of errors (*SSE_D_*) was minimized. We computed the fraction of total variance in the data explained by the fits (*qval*).

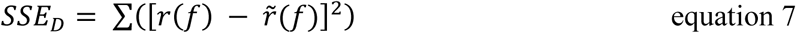

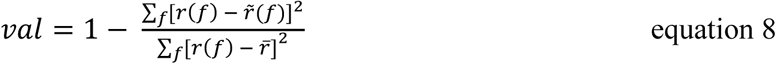

here, 𝑟(𝑓) indicates the observed response and *r̃*(*f*) indicates the fitted value. *r̄* is the mean change in amplitude across all conditions. *qval* value closer to 1 indicates a decent fit.

### Statistical Analysis

To check the statistical significance of the laterality index, we did a right-tailed Wilcoxon signed-rank test, with the alternate hypothesis that LI values were greater than zero, implying right-hemisphere dominance of ASSRs. We also used the right-tailed Wilcoxon signed-rank test to check if masks with modulation frequencies closer to the modulation frequency of the target are more suppressive than masks with modulation frequencies that are farther away. The null hypothesis was that all mask frequencies were equally suppressive.

### Data and Code Availability

Data to generate the figures is available on https://osf.io/b35km/ and all the codes used for this study are available on GitHub (https://github.com/Divya-Gulati/Modulation-Frequency-Masking-).

## Results

We recorded 64-channel EEG data from 24 participants (12 females) while they passively listened to sound stimuli with closed eyes. In the first experiment, only a single amplitude-modulated sinusoid was presented at different modulation frequencies and depths. In the second experiment, two modulation frequencies were concurrently used, with the ‘target’ always modulated at 40Hz, and the ‘mask’ was modulated at frequencies between 22-58Hz (in steps of 6Hz). Both modulation frequencies were independently modulated at different depths. ASSRs evoked in response to stimuli were characterized using their amplitude spectrum, computed using Fourier analysis.

### Auditory Steady State Response scalp topography

Figure 2 shows the subject-averaged ASSR response for an amplitude-modulated sinusoid presented at 100% modulation depth across all the electrodes placed on the scalp. Figure 2A shows the time-frequency spectrograms for the change in power for a stimulus modulated at 40Hz from baseline stimuli, i.e., when no sound was played. We observed a robust ASSR response in the parieto-temporal, parietal, parieto-occipital, and occipital electrodes. Figure 2B shows the change in amplitude at the respective modulation frequency of the sound played. Here, electrodes are arranged in groups (shown in different colours) and in ascending order of increasing response amplitude at 40 Hz for each group. Across different modulation frequencies, we observed a stronger response for the same set of electrodes, i.e., in temporal, parietal, and occipital regions. These results are consistent with prior studies that also obtained responses in similar electrodes when the reference electrode was at the vertex (Gransier et al., 2017, 2021; Vanvooren et al., 2014). Strong responses in parietal and occipital regions could be because of the location of the reference electrode as its position can change the ASSR strength and phase, thereby effectively changing the localization of response (Lu et al., 2022; Sonck et al., 2025-see figures 3 and 6). Based on these results, electrodes were chosen and divided into left and right groups (Figure 2A, bottom right inset; left-blue, right-orange) to allow comparison across hemispheres for further analysis.

**Figure 2.**
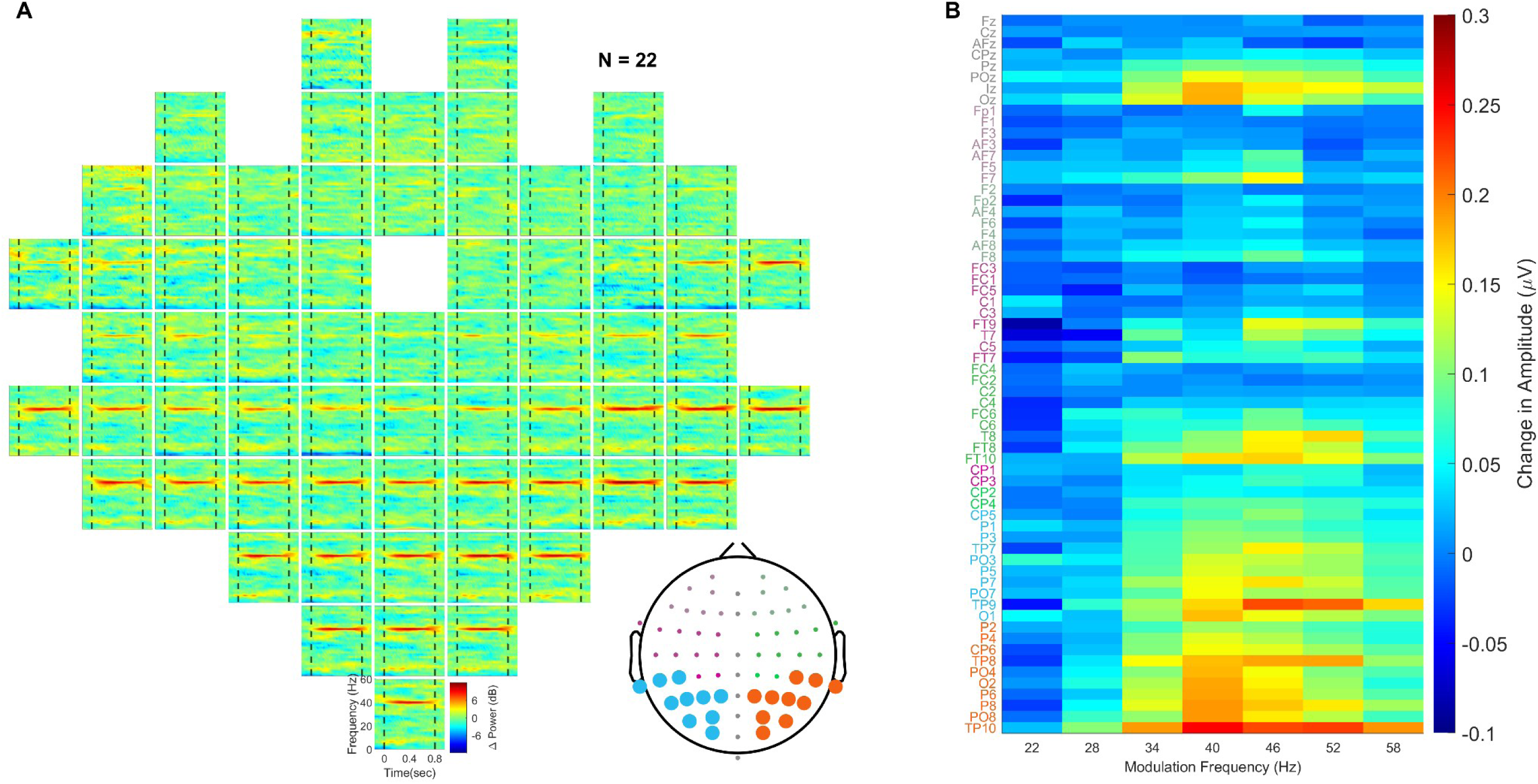
Response across the scalp for amplitude-modulated sinusoids at 100% modulation depth. (A) Time-frequency plots for all 64 channels placed on the scalp show the subject-averaged change in power for a 1kHz sinusoid modulated at 40Hz (N = 22; frequency: 0 - 60Hz, vertical axis and time: -150 to 950ms, horizontal axis). The plots are arranged according to the 64-channel layout (actiCap), with FCz as the reference electrode (empty box in the middle). The dashed lines mark the stimulus presentation duration (0 to 800ms). The colorbar indicates the log power ratio in decibels (dB). The blue and orange color marked electrodes in the right bottom topoplot show the electrode groups of the left and right hemispheres used for further analysis. (B) Subject-averaged change in amplitude during stimulus condition (250 to 750ms) from the baseline (-500 to 0ms) shown as a function of modulation frequency and recording channel. Colorbar indicates change in amplitude in μV at the respective modulation frequency. The electrodes are arranged in order of increasing amplitude at 40Hz for each group. Color of the electrode label indicates the group that electrode belongs to (refer to panel A inset for the group location).

**Figure 3.**
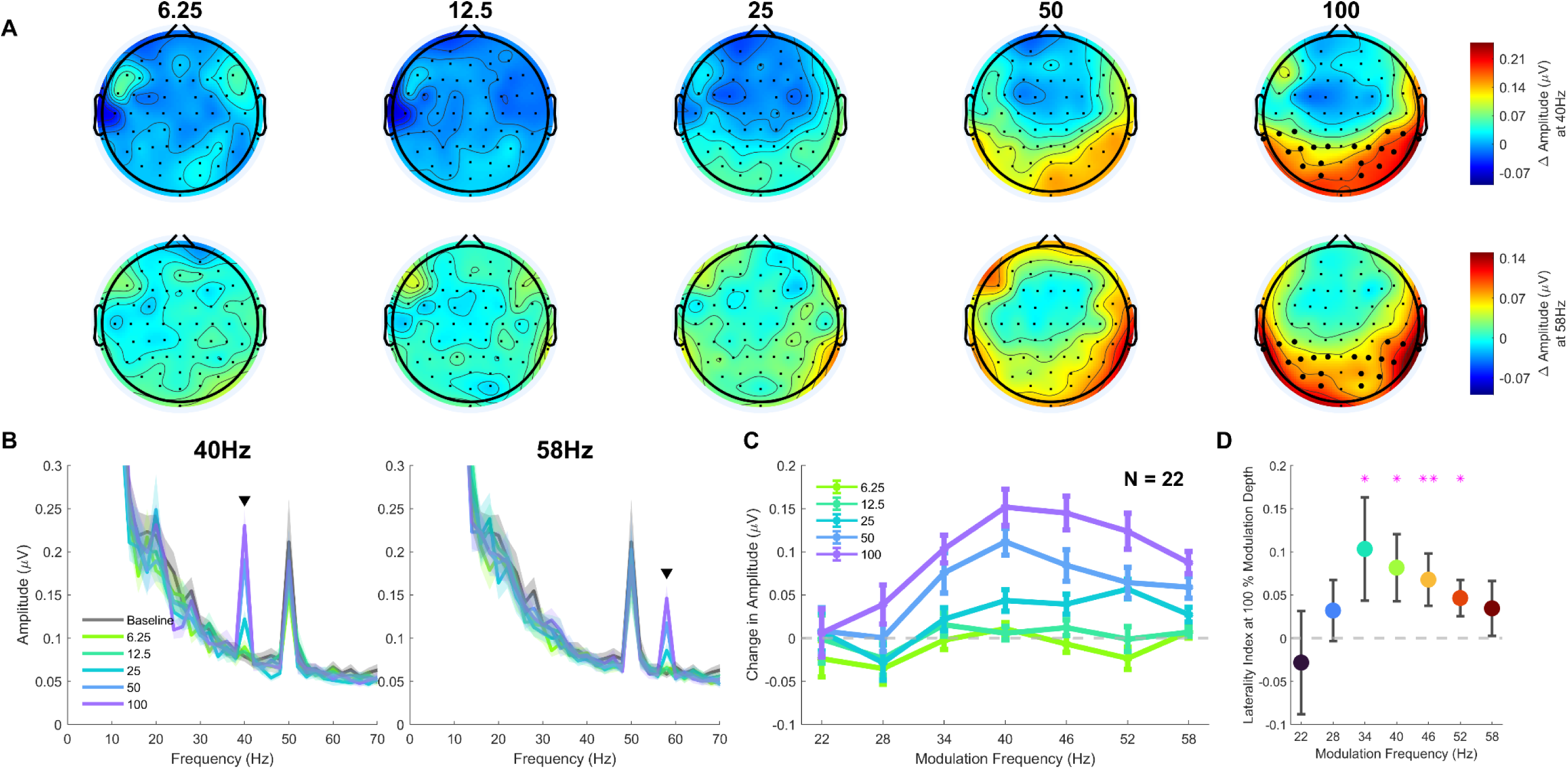
ASSR tuning profiles of amplitude-modulated sinusoids as a function of modulation depth. (A) Scalp maps showing averaged change in amplitude across subjects (N = 22) for 40 (top row) and 58Hz (bottom row) amplitude-modulated sinusoids at five different modulation depths (6.25 to 100% on a log scale). Colorbar indicates change in amplitude in μV at the respective modulation frequency. Electrodes used for analysis are highlighted in bold in the last scalp map in each row. (B) Amplitude spectra averaged across the left and right electrode groups for 40 (left panel) and 58Hz (right panel) amplitude-modulated sinusoids at various modulation depths. (C) ASSR tuning curves show the change in amplitude at the respective modulation frequency of the amplitude-modulated sinusoid as a function of modulation depth, averaged across left and right electrode groups and subjects. (D) Laterality Index values were calculated for amplitude during the stimulus period for amplitude-modulated sounds at different modulation frequencies at 100% modulation depth. The central node represents the median, and the error bar represents the standard error. Star value indicates the p-values for right-tailed Wilcoxon signed rank test (p<0.05: * and p<0.01: **).

### Auditory Steady State Response characteristics

Next, we wanted to characterize the ASSR response for different modulation frequencies as a function of modulation depth. Figure 3 (A-C) shows that a discernible steady state response was observed for modulation depths of 25% and above, which monotonically increased with an increase in modulation depth. In Figure 3B, clear spectral peaks can be observed at the respective modulation frequencies, and a 50Hz line noise artefact is also visible. The strongest ASSR response at 100% modulation depth was observed for 40Hz modulation frequency. In Figure 3A, in the top row, it can be observed that ASSR responses appear to be lateralized, with stronger responses in the right hemisphere. We calculated the laterality index (Figure 3D) to confirm the same. We found lateralization towards the right hemisphere for amplitude modulation sounds with modulation frequency between 34-52 Hz (right-tailed Wilcoxon signed rank test – uncorrected p-value at 34Hz- 0.0489; 40Hz- 0.0189; 46Hz - 0.0043; 52Hz- 0.0174; note that upon Bonferroni correction, the laterality index remained significant only at 46 Hz, although it was positive at all frequencies except 22 Hz). These results align with those reported in previous studies (Gransier et al., 2021; Kuwada et al., 1986; Ross et al., 2005; Weisz & Lithari, 2017).

### Suppression is band-pass: mask modulation frequencies closer to the target modulation frequency suppress strongly

Next, we characterized how the presence of two modulation frequencies in the stimulus can have interactional effects on ASSR amplitudes. To answer this, we measured the ASSR when the carrier sinusoid amplitude was modulated with two sinusoids at varying modulation frequencies. We chose the 40Hz stimulus as the target because we obtained the maximum change in amplitude for this modulation frequency. We then analyzed how the target amplitude at 40Hz changes with the modulation frequency of the mask. Figure 4A depicts the amplitude spectra when both were modulated at 50% depth, averaged separately for left and right electrodes. We observed that the response at 40Hz was reduced when the modulation frequency of the mask differed from that of the target (blue or orange trace) compared to when the target was presented alone at 50% modulation depth (green trace, change in amplitude shown by black bars). A facilitation in the response amplitude was observed when both target and mask modulation frequencies were the same. Suppression was visible in the electrode groups of both hemispheres. We then plotted response amplitudes at 40Hz as a function of modulation frequencies (Figure 4B). We observed that the mask with a modulation frequency closer to that of the target appeared more suppressive than a mask with a modulation frequency farther away. To quantify this, we calculated the amplitude difference between the response amplitude of the mask with different modulation frequencies. We made two such frequency pairs (22-34Hz and 58-46Hz) and averaged the subtracted responses obtained across the pair for each subject (i.e., amplitude difference calculated as (A_22_ – A_34_ )/2 + (A_58_ – A_46_)/2, where A_x_ is the ASSR amplitude at a modulation frequency of x). We observed (Figure 4C) that the amplitude difference was positive (i.e., more suppression at frequencies nearer to 40 Hz than farther away) for both groups of electrodes (Figure 4D), but was more prominent for the right electrode group (right-tailed Wilcoxon signed rank test, p-value: 0.0109). For the left electrode group, the amplitude difference was positive but did not reach significance (right-tailed Wilcoxon signed rank test, p-value: 0.0631), primarily because masks with modulation frequencies lower than 40Hz were equally suppressive (Figure 4B).

**Figure 4.**
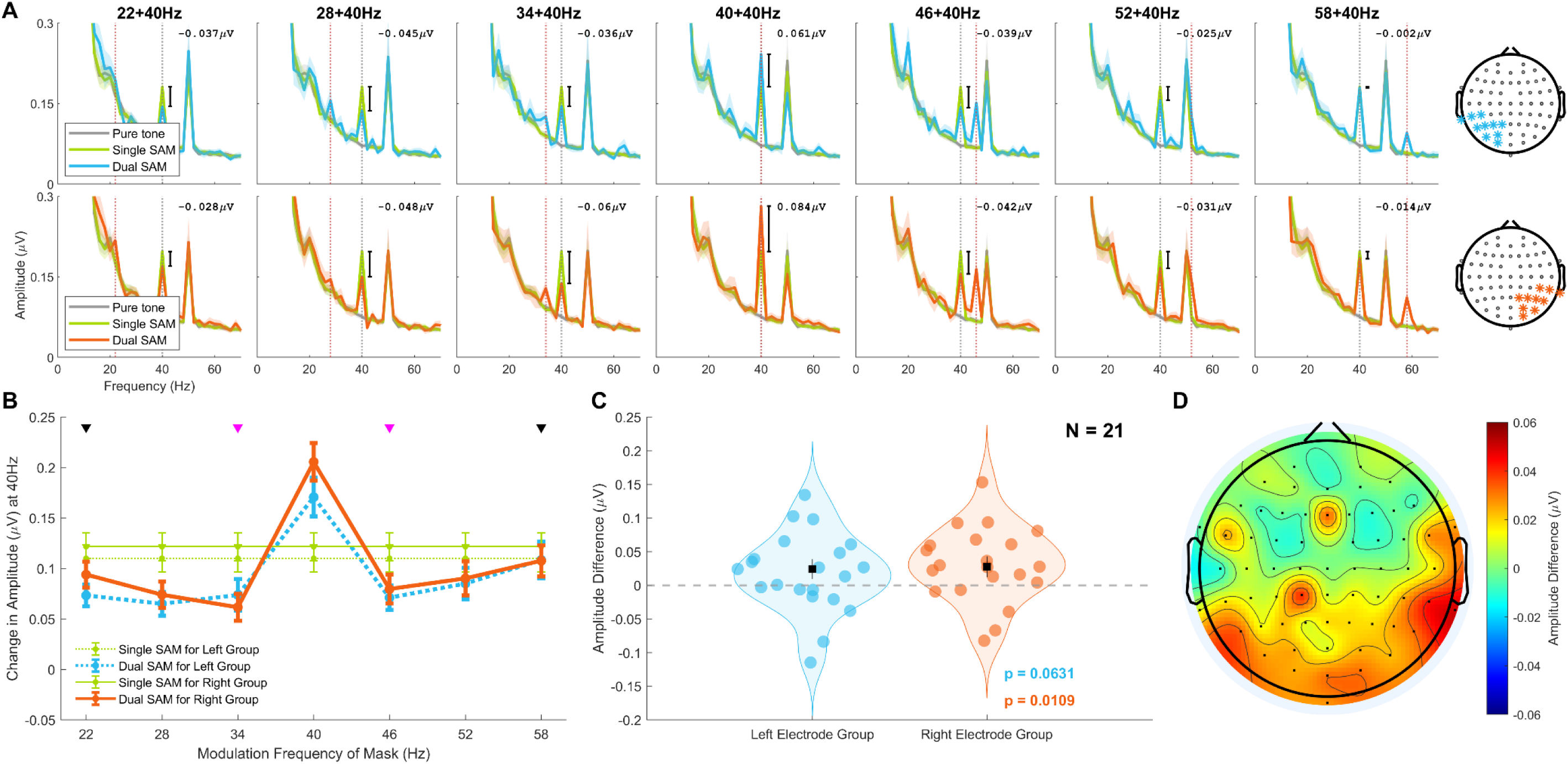
Amplitude response suppression as a function of different modulation frequencies across left and right hemisphere electrode groups. (A) Mean amplitude spectra averaged across subjects (N= 21) for the left (top row) and right (bottom row) groups of electrodes. Each column corresponds to a different modulation frequency of the mask amplitude-modulated sound presented with a target modulation at 40Hz, indicated by the cyan (top row) or orange trace (bottom row). The modulation depth for both target and mask was kept at 50%. The grey trace shows an unmodulated 1kHz pure tone (mostly hidden behind other traces except at 40 Hz), and the green trace shows the amplitude-modulated sinusoid at 40Hz (target ‘only’ condition when modulation depth for mask was at zero; Single SAM); both are the same across all plots across each row. The dotted grey and reddish pink lines indicate the modulation frequency of the target and mask sinusoid, respectively. The numbers indicate the difference in amplitude at 40Hz between the dual SAM and single SAM conditions. (B) Change in amplitude at 40Hz from baseline (1kHz unmodulated pure tone) is shown as a function of the modulation frequency of the mask sinusoid, averaged for the left (dotted lines) and right groups (solid lines) of electrodes under different sound conditions with modulation depth fixed at 50%. Error bars indicate the ±SEM. (C) To determine whether a mask with a modulation frequency closer to that of the target is more suppressive than a mask with a modulation frequency farther from the target, we calculated the amplitude difference between the two conditions across two frequency pairs, as indicated by the arrows (22–34 Hz and 58–46 Hz; black-magenta arrows) in panel B. These differences were averaged for each subject. The violin plots display the distribution of these averaged amplitude differences for the left (blue) and right (orange) electrode groups. The black markers with error bars in the centre represent the median ± standard error (SE). p-values obtained from the right-tailed Wilcoxon signed rank test are indicated on the bottom right. (D) Scalp map showing the amplitude difference averaged across subjects (N = 21) across 64 electrodes. The colour bar indicates amplitude difference in μV.

Additionally, we could observe that for both the electrode groups, the response amplitude when the mask had a modulation frequency of 58Hz was equivalent to when only the target was presented alone, but this was not the case for the mask with a modulation frequency of 22Hz. Even though 22Hz amplitude-modulated sound, when presented alone, did not evoke a strong response (Figure 3C), it was still very suppressive when presented with the target (Figure 4B).

### A slowly varying normalization drive can account for ASSR interaction effects

To explain how ASSR responses at target frequency change as a function of suppression arising from the mask, we used an optimal-tuned normalization model, which has been recently used in the context of SSVEP interactions in the primary visual cortex of macaques (Gulati & Ray, 2025). To estimate the model parameters, we independently modulated the target and mask modulation frequency at three different depths (0, 25, and 50%). We therefore obtained nine response profiles, one for each combination of target and mask modulation depth (similar to one profile shown in Figure 4B, when both mask and target were at 50% modulation depths). From each profile, we obtained 6 data points (the response at 40 Hz, which showed a facilitation, was not considered), or a total of 54 data points, which were fitted using a 6-parameter model described in equations 5 and 6. Figure 5A shows the subject-averaged change in amplitude at 40Hz as a function of target and mask modulation depth and frequency (indicated by filled circles). This data was averaged across electrode groups of both right and left hemispheres for each subject, as we did not observe any response suppression lateralization (Figure 4D). The response suppression was stronger when mask and target modulation depths were increased; the overall trend across different depths remained the same. We averaged this data across selected electrodes across both hemispheres for each subject and then used the optimal-tuned normalization model to fit each subject’s data. Fit for subjects where the model could capture explained variance of more than 40% were averaged (14 out of 21 subjects) and plotted in Figure 5 (dashed lines). On average, the model captured 62% variance in the data. Similar to our previous result with SSVEPs (Gulati & Ray, 2025), we obtained a band-pass shaped, target frequency-dependent suppression profile (Figure 5B) with a subject-averaged low-pass filter cutoff of ∼7.5Hz. Median free parameters for the model are shown in Table 1.

**Figure 5.**
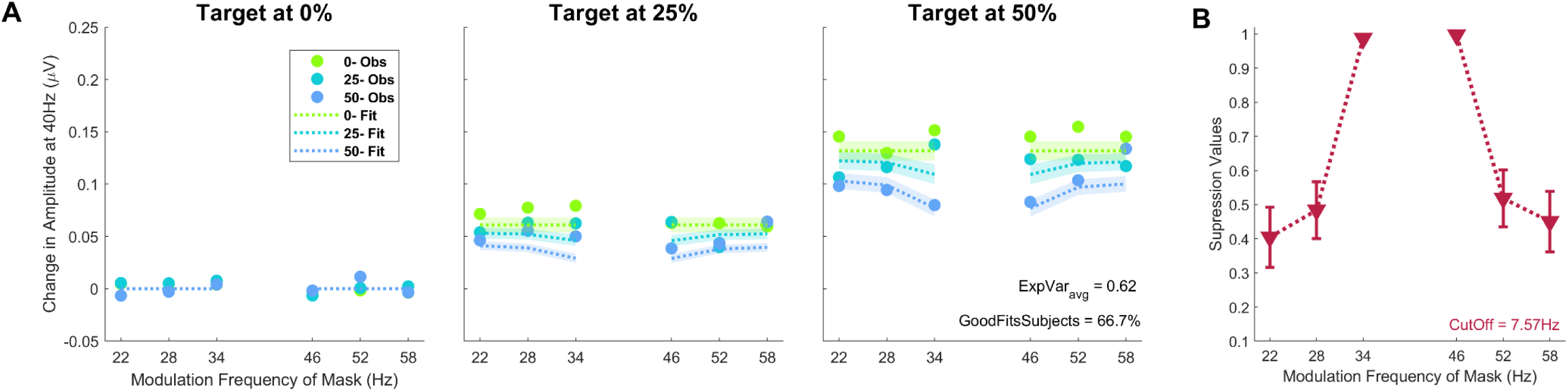
Amplitude response suppression as a function of target and mask modulation depth and frequency captured by an optimal-tuned normalization model. (A) Change in amplitude at 40Hz from baseline condition (1kHz unmodulated pure tone) averaged across electrode groups and subjects, with target and mask amplitude-modulated sounds presented at various modulation depths. Each sound can take one of the three modulation depth values from 0, 25% or 50%. Each panel represents the modulation depth of the target sinusoid, and within each panel, three different dot colours represent the modulation depth of the mask sinusoid (green to blue indicating increasing depth). Dotted lines with shaded regions indicate the model fits averaged across subjects (mean ± SEM) with explained variance >= 0.4. The percentage of subjects with explained variance >= 0.4 is represented as ‘GoodFitsSubjects’, and ‘ExpVar_avg_’ gives the averaged explained variance. (B) Averaged suppression values obtained from the model as a function of the modulation frequency of the mask. The profile for each subject was calculated from the free parameter values obtained from the model using equation 6 and was normalized before averaging across subjects. The averaged low-pass filter cutoff obtained across subjects from the model is represented by ‘CutOff’.

**Table 1:** Showing median ± bootstrapped SEM values for parameters obtained from the optimal-tuned normalization model.

| *Parameters* | *Obtained Value* |
| --- | --- |
|  | *(Median ± bootstrapped SEM)* |
| $\sigma$ | 0.4472 ± 0.0528 |
| $L_{amp}$ | 0.5003 ± 0.0105 |
| *Cutoff* | 7.0567 ± 0.1601 |
| $\alpha_1$ | 3.0684 ± 0.0107 |
| $\alpha_2$ | 2.2246e-09 ± 5.4932e-07 |
| $ScF$ | 0.8031 ± 0.0284 |

## Discussion

We systematically captured the interaction between ASSRs and their underlying mechanisms when two SAM stimuli are presented together. In our experiment 1, we presented a single SAM stimulus of various modulation frequencies and depths to quantify the ASSR response in the EEG. We observed the strongest response at 40Hz and saw a monotonic increase in ASSR amplitude across all modulation frequencies as a function of modulation depth, similar to that observed previously (Galambos et al., 1981; Ross et al., 2000). In experiment 2, we simultaneously modulated a pure tone at two different modulation frequencies. We showed evidence for the dependence of the amount of modulation masking of ASSRs on the temporal characteristics of stimuli. On this data, we implemented a slow-varying normalization model that low-pass filters the normalization drive. The model architecture used here was originally used to explain SSVEP responses (Gulati & Ray, 2025). The same model also explained ASSR interactions, suggesting shared operations across the two sensory modalities.

### Comparison with previous studies

Previous MEG studies (Kaneko et al., 2003; Kawase et al., 2012) have observed lateralization of ASSR suppression due to the dichotic presentation of stimuli (i.e., target and mask stimuli presented separately in each ear). However, we did not observe distinct lateralization of ASSR suppression, although it was marginally more pronounced towards the right hemisphere.

While psychoacoustic studies (Bacon & Grantham, 1989; Grantham & Bacon, 1991; Sek et al., 2015) have shown that masking profile is dependent on the modulation frequency of the mask stimulus, a study examining (Gransier et al., 2017) the dichotic interactions between ASSRs generated between 30-50Hz did not find this dependence. Our results, however, did reveal dependence of the modulation frequency of the mask stimulus on ASSR suppression. We obtained a band-pass shape suppression profile different from that obtained perceptually (Bacon & Grantham, 1989; Grantham & Bacon, 1991; Sek et al., 2015).

These discrepancies between our results and those reported previously could be attributed to minor differences in the stimulus design and to diotic stimulus presentation (i.e., presentation of both target and mask stimuli in the same ear) in our case. ASSR attenuation is observed to be dependent on the carrier frequencies of the two SAM stimuli and only occurs when the sound intensity is high (at 75dB SPL) (John et al., 1998). This is potentially because the tuning of modulation filters depends on the carrier frequency and level of the SAM stimulus (Simpson et al., 2013). In our study, we fixed the intensity and used a single carrier frequency to eliminate their influence.

### Normalization model architecture

In the slow-varying normalization model (Gulati & Ray, 2025), masking normalization depends on the orientation selectivity of the V1 neurons, which act as different filters. In the auditory domain, several temporal modulation detection studies (Bacon & Grantham, 1989; Houtgast, 1989; Yost et al., 1989) have pointed to the existence of modulation-frequency-specific channels at the central auditory system level, referred to as modulation filterbanks (Figure 1). As shown in Figure 1, previous models (Dau et al., 1997a, 1997b; Yost et al., 1989) proposed for amplitude-modulated sound perception suggest that demodulation starts in the periphery itself. The following stages consist of hierarchical band-limited modulation filters, decomposing the SAM envelope into modulation components. This means that each processing stage in the auditory pathway leading up to the auditory cortex preferentially extracts a given modulation frequency range, with each stage processing slower modulations than the last, as confirmed by fMRI (Giraud et al., 2000) and sEEG (Liégeois-Chauvel et al., 2004) studies in humans. These studies showed that the human auditory cortex can respond well to modulations slower than ∼60 Hz. To keep the model comparable to the previous SSVEP model and since ASSR interactions below 60Hz are cortically originated (Kuwada et al., 1986), we only considered the modulation filter bank at the level of the auditory cortex. The modulation filters essentially represent the spectro-temporal receptive field (STRF) of the auditory neurons (Chi et al., 2005). It has been previously shown that STRF measured for the envelope of the stimuli alone is equivalent to the STRF measured for the complete stimulus (Elhilali et al., 2004), indicating that primary auditory cortex neurons can precisely follow the envelope of the stimuli at a particular carrier frequency, and thus giving a strong response at the modulation frequency for SAM stimuli.

Although the normalization architecture is similar across visual and auditory modalities, differences in filter characteristics lead to different predictions. In the case of vision, the filters are sensitive to orientation but independent of modulation frequency, and therefore signals pass through the same filter if the stimulus orientations are similar, irrespective of the SSVEP tagging frequencies used. However, in the case of the auditory system, the filters themselves are sensitive to the modulation frequency. So the signals should pass through the same filter for nearby ASSR frequencies but through different filters when they are far apart, leading to a mixture of band-pass or low-pass behaviour for nearby versus far-away ASSR frequencies.

In our model, to account for these differences, we kept both within-filter and across-filter summation, with scaling factors of *α_1_* and *α_2_*, respectively (see equation 6). However, *α_2_* was practically zero in the fitted data (Table 1), suggesting that indeed both ASSRs were passing through the same filter. This is consistent with prior psychoacoustical (Houtgast, 1989) and modelling studies (Dau et al., 1997a; Ewert & Dau, 2000; Joosten et al., 2016) that have observed that the filter bandwidth of modulation filters is ½-1 octave wide around the centre frequency and the Q-values, which indicate the ratio of center frequency to the bandwidth of the filter, lie between 1 and 2. So, since the modulation frequencies we used in the current study all lay between one octave of the target frequency and, therefore, could have passed through the same filters, thus giving us a band-pass suppression profile. But the suppression profile could be very different if the target frequency was lower (say 10 Hz), where the modulation filterbanks are modelled to have a bandwidth of only ∼5 Hz (Dau et al., 1997a).

When two modulation frequencies pass through the non-linear stage after summing up, intermodulation terms at sums and differences of modulation frequencies in the amplitude spectra could also be observed. This presence of IM terms validates that the same filter processes modulation frequencies. Previous reports have shown the presence of intermodulation terms (Draganova et al., 2002; Xiang et al., 2013) when concurrent SAM stimuli were played. We did observe a peak at the *f_1_ + f_2_* intermodulation term, but only for the mask at 46Hz. Absence of peaks for other modulation frequencies could be attributed to low ASSR amplitude at all the other respective mask modulation frequencies, and IM peaks are even weaker than them by comparison. Since subjects had eyes closed during the baseline and stimulus periods, they had strong alpha oscillations in the range of 6-14Hz. And for the stimuli we used, most *f_1_–f_2_* intermodulation terms lay within that range and therefore, could not be observed.

Several studies that have modelled processing of temporally modulated stimuli by neurons in the auditory pathway have considered the inhibitory signal in the model to be slower in comparison to the excitatory signals (Eggermont, 1999; Elhilali et al., 2004; Nelson & Carney, 2007), which is in agreement with our slow-varying normalization drive.

### Similarities and differences with visual modality

The band-pass suppression profile for ASSRs we obtained is similar to that obtained when the component gratings of the counterphase plaid stimuli were parallel. In our study, we obtained the filter cutoff of the low-pass filter to be around ∼7.5Hz, which matches the cutoffs obtained for parallel plaids, which were between 7-11Hz (Gulati & Ray, 2025). This corresponds to an integration time (1/2π*f*) of about ∼21ms, which is in agreement with the previous study that found a temporal filter delay of ∼10-15ms was best for capturing the responses of primary and secondary auditory cortex neurons of cats towards periodic click stimuli (Eggermont, 1999). GABA_A_ is found to be involved in masking-induced suppression of SSVEP responses (Morrone et al., 1987) and the generation of 40Hz ASSRs (Toso et al., 2024). This alludes that GABA_A_ might also be involved in ASSR suppression. Parvalbumin neurons, which are involved in divisive scaling of responses in the visual cortex (Atallah et al., 2012), are not found to be the main contributors in the auditory cortex (Cooke et al., 2020). Instead, neuromodulatory zinc signalling has been found to facilitate cortical gain control (Cody & Tzounopoulos, 2022). These findings suggest that the auditory and visual cortex might use different underlying mechanisms to implement the same algorithm.

### Study Limitations

We note that the model could fit responses for only 14 out of 21 good subjects; this could be because the ASSR tuning itself was observed to vary across participants, similar to that observed previously (Gransier et al., 2021). We acknowledge the need to consider our results for suppression profile shape and filter cutoff with caution, as our stimulus set was limited. We only used seven mask modulation frequencies, each at a gap of 6Hz, with the target always fixed at 40Hz. Therefore, unlike previous studies with gratings, where 15 mask temporal frequencies, separated by 2Hz, were used (Gulati & Ray, 2025), the parameter space used to obtain the suppression profile in the current study was not densely sampled. We limited the number of modulation frequencies used to not lengthen the recording time further, as it was already ∼3 hours per subject. Therefore, further studies with fine sampling of mask modulation frequencies, at different target frequencies, are needed to characterize these ASSR interactions comprehensively.

## Acknowledgements

This work was supported by Wellcome Trust/DBT India Alliance (Senior fellowship IA/S/18/2/504003 to S.R.), DBT-IISc Partnership Programme and Pratiksha Trust. The Council of Scientific and Industrial Research (CSIR) awarded D.G. a senior research fellowship.

## References

1. Atallah, B. V., Bruns, W., Carandini, M., & Scanziani, M. (2012). Parvalbumin-Expressing Interneurons Linearly Transform Cortical Responses to Visual Stimuli. Neuron, 73(1), 159–170. 10.1016/j.neuron.2011.12.013

2. Bacon, S. P., & Grantham, D. W. (1989). Modulation masking: Effects of modulation frequency, depth, and phase. The Journal of the Acoustical Society of America, 85(6), 2575– 2580. 10.1121/1.397751

3. Bokil, H., Andrews, P., Kulkarni, J. E., Mehta, S., & Mitra, P. P. (2010). Chronux: A platform for analyzing neural signals. Journal of Neuroscience Methods, 192(1), 146–151. 10.1016/j.jneumeth.2010.06.020

4. Candy, T. R., Skoczenski, A. M., & Norcia, A. M. (2001). Normalization models applied to orientation masking in the human infant. The Journal of Neuroscience: The Official Journal of the Society for Neuroscience, 21(12), 4530–4541. 10.1523/JNEUROSCI.21-12-04530.2001

5. Chi, T., Ru, P., & Shamma, S. A. (2005). Multiresolution spectrotemporal analysis of complex sounds. The Journal of the Acoustical Society of America, 118(2), 887–906. 10.1121/1.1945807

6. Cody, P. A., & Tzounopoulos, T. (2022). Neuromodulatory Mechanisms Underlying Contrast Gain Control in Mouse Auditory Cortex. Journal of Neuroscience, 42(28), 5564–5579. 10.1523/JNEUROSCI.2054-21.2022

7. Cooke, J. E., Kahn, M. C., Mann, E. O., King, A. J., Schnupp, J. W. H., & Willmore, B. D. B. (2020). Contrast gain control occurs independently of both parvalbumin-positive interneuron activity and shunting inhibition in auditory cortex. Journal of Neurophysiology, 123(4), 1536–1551. 10.1152/jn.00587.2019

8. Cooke, J. E., King, A. J., Willmore, B. D. B., & Schnupp, J. W. H. (2018). Contrast gain control in mouse auditory cortex. Journal of Neurophysiology, 120(4), 1872–1884. 10.1152/jn.00847.2017

9. Dau, T., Kollmeier, B., & Kohlrausch, A. (1997a). Modeling auditory processing of amplitude modulation. I. Detection and masking with narrow-band carriers. The Journal of the Acoustical Society of America, 102(5), 2892–2905. 10.1121/1.420344

10. Dau, T., Kollmeier, B., & Kohlrausch, A. (1997b). Modeling auditory processing of amplitude modulation. II. Spectral and temporal integration. The Journal of the Acoustical Society of America, 102(5), 2906–2919. 10.1121/1.420345

11. Delorme, A., & Makeig, S. (2004). EEGLAB: An open source toolbox for analysis of single-trial EEG dynamics including independent component analysis. Journal of Neuroscience Methods, 134(1), 9–21. 10.1016/j.jneumeth.2003.10.009

12. Ding, J., Sperling, G., & Srinivasan, R. (2006). Attentional modulation of SSVEP power depends on the network tagged by the flicker frequency. Cerebral Cortex (New York, N.Y. : 1991), 16(7), 1016–1029. 10.1093/cercor/bhj044

13. Draganova, R., Ross, B., Borgmann, C., & Pantev, C. (2002). Auditory Cortical Response Patterns to Multiple Rhythms of AM Sound. Ear and Hearing, 23(3), 254.

14. Eggermont, J. J. (1999). The Magnitude and Phase of Temporal Modulation Transfer Functions in Cat Auditory Cortex. The Journal of Neuroscience, 19(7), 2780–2788. 10.1523/JNEUROSCI.19-07-02780.1999

15. Elhilali, M., Fritz, J. B., Klein, D. J., Simon, J. Z., & Shamma, S. A. (2004). Dynamics of precise spike timing in primary auditory cortex. The Journal of Neuroscience: The Official Journal of the Society for Neuroscience, 24(5), 1159–1172. 10.1523/JNEUROSCI.3825-03.2004

16. Ewert, S. D., & Dau, T. (2000). Characterizing frequency selectivity for envelope fluctuations. The Journal of the Acoustical Society of America, 108(3), 1181–1196. 10.1121/1.1288665

17. Fujiki, N., Jousmäki, V., & Hari, R. (2002). Neuromagnetic Responses to Frequency-Tagged Sounds: A New Method to Follow Inputs from Each Ear to the Human Auditory Cortex during Binaural Hearing. Journal of Neuroscience, 22(3), RC205–RC205. 10.1523/JNEUROSCI.22-03-j0003.2002

18. Galambos, R., Makeig, S., & Talmachoff, P. J. (1981). A 40-Hz auditory potential recorded from the human scalp. Proceedings of the National Academy of Sciences, 78(4), 2643–2647. 10.1073/pnas.78.4.2643

19. Giraud, A.-L., Lorenzi, C., Ashburner, J., Wable, J., Johnsrude, I., Frackowiak, R., & Kleinschmidt, A. (2000). Representation of the Temporal Envelope of Sounds in the Human Brain. Journal of Neurophysiology, 84(3), 1588–1598. 10.1152/jn.2000.84.3.1588

20. Gransier, R., Hofmann, M., van Wieringen, A., & Wouters, J. (2021). Stimulus-evoked phase-locked activity along the human auditory pathway strongly varies across individuals. Scientific Reports, 11(1), 143. 10.1038/s41598-020-80229-w

21. Gransier, R., van Wieringen, A., & Wouters, J. (2017). Binaural Interaction Effects of 30–50 Hz Auditory Steady State Responses. Ear and Hearing, 38(5), e305. 10.1097/AUD.0000000000000429

22. Grantham, D. W., & Bacon, S. P. (1991). Binaural modulation masking. The Journal of the Acoustical Society of America, 89(3), 1340–1349. 10.1121/1.400657

23. Gulati, D., & Ray, S. (2024). Auditory and Visual Gratings Elicit Distinct Gamma Responses. eNeuro, 11(4), ENEURO.0116-24.2024. 10.1523/ENEURO.0116-24.2024

24. Gulati, D., & Ray, S. (2025). Slow-varying normalization explains diverse temporal frequency masking interactions in the macaque primary visual cortex (p. 2025.03.11.642541). bioRxiv. 10.1101/2025.03.11.642541

25. Houtgast, T. (1989). Frequency selectivity in amplitude-modulation detection. The Journal of the Acoustical Society of America, 85(4), 1676–1680. 10.1121/1.397956

26. Ishida, I. M., & Stapells, D. R. (2012). Multiple-ASSR Interactions in Adults with Sensorineural Hearing Loss. International Journal of Otolaryngology, 2012, 802715. 10.1155/2012/802715

27. John, M. S., Lins, O. G., Boucher, B. L., & Picton, T. W. (1998). Multiple auditory steady-state responses (MASTER): Stimulus and recording parameters. Audiology: Official Organ of the International Society of Audiology, 37(2), 59–82. 10.3109/00206099809072962

28. Joosten, E. R. M., Shamma, S. A., Lorenzi, C., & Neri, P. (2016). Dynamic Reweighting of Auditory Modulation Filters. PLOS Computational Biology, 12(7), e1005019. 10.1371/journal.pcbi.1005019

29. Kaneko, K., Fujiki, N., & Hari, R. (2003). Binaural interaction in the human auditory cortex revealed by neuromagnetic frequency tagging: No effect of stimulus intensity. Hearing Research, 183(1), 1–6. 10.1016/S0378-5955(03)00186-2

30. Kawase, T., Maki, A., Kanno, A., Nakasato, N., Sato, M., & Kobayashi, T. (2012). Contralateral white noise attenuates 40-Hz auditory steady-state fields but not N100m in auditory evoked fields. NeuroImage, 59(2), 1037–1042. 10.1016/j.neuroimage.2011.08.108

31. King, A. J., & Nelken, I. (2009). Unraveling the principles of auditory cortical processing: Can we learn from the visual system? Nature Neuroscience, 12(6), 698–701. 10.1038/nn.2308

32. Kuwada, S., Batra, R., & Maher, V. L. (1986). Scalp potentials of normal and hearing-impaired subjects in response to sinusoidally amplitude-modulated tones. Hearing Research, 21(2), 179–192. 10.1016/0378-5955(86)90038-9

33. Lazzouni, L., Ross, B., Voss, P., & Lepore, F. (2010). Neuromagnetic auditory steady-state responses to amplitude modulated sounds following dichotic or monaural presentation. Clinical Neurophysiology, 121(2), 200–207. 10.1016/j.clinph.2009.11.004

34. Liégeois-Chauvel, C., Lorenzi, C., Trébuchon, A., Régis, J., & Chauvel, P. (2004). Temporal Envelope Processing in the Human Left and Right Auditory Cortices. Cerebral Cortex, 14(7), 731–740. 10.1093/cercor/bhh033

35. Lins, O. G., Picton, P. E., Picton, T. W., Champagne, S. C., & Durieux-Smith, A. (1995). Auditory steady-state responses to tones amplitude-modulated at 80–110 Hz. The Journal of the Acoustical Society of America, 97(5), 3051–3063. 10.1121/1.411869

36. Lins, O. G., & Picton, T. W. (1995). Auditory steady-state responses to multiple simultaneous stimuli. Electroencephalography and Clinical Neurophysiology/Evoked Potentials Section, 96(5), 420–432. 10.1016/0168-5597(95)00048-W

37. Liza, K., & Ray, S. (2022). Local Interactions between Steady-State Visually Evoked Potentials at Nearby Flickering Frequencies. The Journal of Neuroscience, 42(19), 3965–3974. 10.1523/JNEUROSCI.0180-22.2022

38. Lopez, M.-A., Pomares, H., Pelayo, F., Urquiza, J., & Perez, J. (2009). Evidences of cognitive effects over auditory steady-state responses by means of artificial neural networks and its use in brain–computer interfaces. Neurocomputing, 72(16), 3617–3623. 10.1016/j.neucom.2009.04.021

39. Lu, H., Mehta, A. H., & Oxenham, A. J. (2022). Methodological considerations when measuring and analyzing auditory steady-state responses with multi-channel EEG. Current Research in Neurobiology, 3, 100061. 10.1016/j.crneur.2022.100061

40. Morrone, M. C., Burr, D. C., & Speed, H. D. (1987). Cross-orientation inhibition in cat is GABA mediated. Experimental Brain Research, 67(3), 635–644. 10.1007/BF00247294

41. Müller, M. M., Andersen, S., Trujillo, N. J., Valdés-Sosa, P., Malinowski, P., & Hillyard, S. A. (2006). Feature-selective attention enhances color signals in early visual areas of the human brain. Proceedings of the National Academy of Sciences of the United States of America, 103(38), 14250–14254. 10.1073/pnas.0606668103

42. Müller, N., Schlee, W., Hartmann, T., Lorenz, I., & Weisz, N. (2009). Top-Down Modulation of the Auditory Steady-State Response in a Task-Switch Paradigm. Frontiers in Human Neuroscience, 3, 1. 10.3389/neuro.09.001.2009

43. Murty, D. V. P. S., & Ray, S. (2022). Stimulus-induced Robust Narrow-band Gamma Oscillations in Human EEG Using Cartesian Gratings. Bio-Protocol, 12(7), e4379. 10.21769/BioProtoc.4379

44. Nelson, P. C., & Carney, L. H. (2007). Neural Rate and Timing Cues for Detection and Discrimination of Amplitude-Modulated Tones in the Awake Rabbit Inferior Colliculus. Journal of Neurophysiology, 97(1), 522–539. 10.1152/jn.00776.2006

45. Rabinowitz, N. C., Willmore, B. D. B., Schnupp, J. W. H., & King, A. J. (2011). Contrast Gain Control in Auditory Cortex. Neuron, 70(6), 1178–1191. 10.1016/j.neuron.2011.04.030

46. Ross, B., Borgmann, C., Draganova, R., Roberts, L. E., & Pantev, C. (2000). A high-precision magnetoencephalographic study of human auditory steady-state responses to amplitude-modulated tones. The Journal of the Acoustical Society of America, 108(2), 679–691. 10.1121/1.429600

47. Ross, B., Draganova, R., Picton, T. W., & Pantev, C. (2003). Frequency specificity of 40-Hz auditory steady-state responses. Hearing Research, 186(1–2), 57–68. 10.1016/s0378-5955(03)00299-5

48. Ross, B., Herdman, A. T., & Pantev, C. (2005). Right hemispheric laterality of human 40 Hz auditory steady-state responses. Cerebral Cortex (New York, N.Y.: 1991), 15(12), 2029–2039. 10.1093/cercor/bhi078

49. Salelkar, S., & Ray, S. (2020). Interaction between steady-state visually evoked potentials at nearby flicker frequencies. Scientific Reports, 10(1), 5344. 10.1038/s41598-020-62180-y

50. Sek, A., Baer, T., Crinnion, W., Springgay, A., & Moore, B. C. J. (2015). Modulation masking within and across carriers for subjects with normal and impaired hearing. The Journal of the Acoustical Society of America, 138(2), 1143–1153. 10.1121/1.4928135

51. Simpson, A. J. R., Reiss, J. D., & McAlpine, D. (2013). Tuning of Human Modulation Filters Is Carrier-Frequency Dependent. PLOS ONE, 8(8), e73590. 10.1371/journal.pone.0073590

52. Slugocki, C., Bosnyak, D., & Trainor, L. J. (2017). Simultaneously-evoked auditory potentials (SEAP): A new method for concurrent measurement of cortical and subcortical auditory-evoked activity. Hearing Research, 345, 30–42. 10.1016/j.heares.2016.12.014

53. Sonck, R., Vanthornhout, J., Bonin, E., & Francart, T. (2025). Auditory Steady-State Responses: Multiplexed Amplitude Modulation Frequencies to Reduce Recording Time. Ear and Hearing, 46(1), 24. 10.1097/AUD.0000000000001552

54. Toso, A., Wermuth, A. P., Arazi, A., Braun, A., Jong, T. G.-‘t, Uhlhaas, P. J., & Donner, T. H. (2024). 40 Hz Steady-State Response in Human Auditory Cortex Is Shaped by Gabaergic Neuronal Inhibition. Journal of Neuroscience, 44(24). 10.1523/JNEUROSCI.2029-23.2024

55. Tsai, J. J., Wade, A. R., & Norcia, A. M. (2012). Dynamics of normalization underlying masking in human visual cortex. The Journal of Neuroscience: The Official Journal of the Society for Neuroscience, 32(8), 2783–2789. 10.1523/JNEUROSCI.4485-11.2012

56. Vanvooren, S., Poelmans, H., Hofmann, M., Ghesquière, P., & Wouters, J. (2014). Hemispheric Asymmetry in Auditory Processing of Speech Envelope Modulations in Prereading Children. The Journal of Neuroscience, 34(4), 1523–1529. 10.1523/JNEUROSCI.3209-13.2014

57. Vialatte, F.-B., Maurice, M., Dauwels, J., & Cichocki, A. (2010). Steady-state visually evoked potentials: Focus on essential paradigms and future perspectives. Progress in Neurobiology, 90(4), 418–438. 10.1016/j.pneurobio.2009.11.005

58. Weisz, N., & Lithari, C. (2017). Amplitude modulation rate dependent topographic organization of the auditory steady-state response in human auditory cortex. Hearing Research, 354, 102–108. 10.1016/j.heares.2017.09.003

59. Xiang, J., Poeppel, D., & Simon, J. Z. (2013). Physiological evidence for auditory modulation filterbanks: Cortical responses to concurrent modulations. The Journal of the Acoustical Society of America, 133(1), EL7–EL12. 10.1121/1.4769400

60. Yost, W. A., Sheft, S., & Opie, J. (1989). Modulation interference in detection and discrimination of amplitude modulation. The Journal of the Acoustical Society of America, 86(6), 2138–2147. 10.1121/1.398474

